# Identification of novel fructo-oligosaccharide bacterial consumers by pulse metatranscriptomics in a human stool sample

**DOI:** 10.1101/2024.07.31.606081

**Authors:** Catherine Prattico, Emmanuel Gonzalez, Lharbi Dridi, Shiva Jazestani, Kristin E. Low, D. Wade Abbott, Corinne F. Maurice, Bastien Castagner

## Abstract

Dietary fibres influence the composition of the human gut microbiota and directly contribute to its downstream effects on host health. As more research supports the use of glycans as prebiotics for therapeutic applications, the need to identify the gut bacteria that metabolize glycans of interest increases. Fructooligosaccharide (FOS) is a common diet-derived glycan that is fermented by the gut microbiota and has been used as a prebiotic. Despite being well studied, we do not yet have a complete picture of all FOS-consuming gut bacterial taxa. To identify new bacterial consumers, we used a short exposure of microbial communities in a stool sample to FOS or galactomannan as the sole carbon source to induce glycan metabolism genes. We then performed metatranscriptomics, paired with whole metagenomic sequencing (WMS), and 16S amplicon sequencing. The short incubation induced genes involved in carbohydrate metabolism, like carbohydrate-active enzymes (CAZymes), including glycoside hydrolase family 32 genes, which hydrolyze fructan polysaccharides like FOS and inulin. Interestingly, FOS metabolism transcripts were notably overexpressed in *Blautia* species not previously reported to be fructan consumers. We therefore validated the ability of different *Blautia* species to ferment fructans by monitoring their growth and fermentation in defined media. This pulse metatranscriptomics approach is a useful method to find novel consumers of prebiotics and increase our understanding of prebiotic metabolism by CAZymes in the gut microbiota.

**Significance:** Complex carbohydrates are key contributors to the composition of the human gut microbiota and play an essential role in the microbiota’s effects on host health. Understanding which bacteria consume complex carbohydrates, or glycans, provides a mechanistic link between dietary prebiotics and their beneficial health effects, an essential step for their therapeutic application. Here, we used a pulse metatranscriptomics pipeline to identify bacterial consumers based on glycan metabolism induction in a human stool sample. We identified novel consumers of FOS among *Blautia* species, expanding our understanding of this well-known glycan. Our approach can be applied to identify consumers of understudied glycans and expand our prebiotic repertoire. It can also be used to study prebiotic glycans directly in stool samples in distinct patient populations to help delineate the prebiotic mechanism.

## Introduction

The human gut microbiota, composed of the bacterial communities residing in the gastrointestinal tract, plays essential roles in immunological homeostasis, pathogen colonization resistance, and the extraction of energy and nutrients from diet (1–5). These interactions with the host, crucial for health, are largely mediated by the diverse metabolic products synthesized by gut bacteria (6, 7). In particular, complex carbohydrates, or glycans, that are present in dietary fibre serve as an important carbon source for gut bacteria and shape gut microbiota diversity and composition (8–12). The main bacterial metabolic output from glycan metabolism are short chain fatty acids (SCFAs), small molecules with potent immunomodulatory effects on the host (13–15). Thus, dietary glycans can act as prebiotics, substrates selectively metabolized by microorganisms conferring health benefits, in part by promoting the production of SCFAs (14–16).

Fructans have been investigated clinically as a prebiotic for various conditions such as cardiovascular disease, diabetes, and colorectal cancer (17–19). Fructans, found in common dietary sources such as onion and chicory, are fructose polymers with varying degree of polymerization (DP). Levans are β2-6 linked fructans which are mostly produced by bacteria (DP 100 to >1000) and can be found in fermented foods or as thickeners and sweeteners (20). Levans are also produced by plants like timothy grass with a DP of 3-73 but are less commonly found in the diet (21). Fructo-oligosaccharides (FOSs) are short-chain β2-1 linked fructans with 1-10 DP, while inulin has a longer chain length of 10-100 DP (22–25). Inulin has been shown to prevent anastomotic leaks from colorectal cancer and promote response to cancer immunotherapy in mouse models and is therefore currently undergoing clinical trials in these conditions (NCT05860322, NCT05821751), as well as others (19, 26–29). However, these same fructan prebiotics can have negative effects in other diseases (30, 31). For example, incomplete inulin degradation can promote inflammation in inflammatory bowel disease patients who lack fermentative microbial activities (32). As bacteria metabolize prebiotics in a species- and sometimes strain-specific manner, the composition of the gut microbiota can determine distinct metabolite production (33, 34). Moreover, because different individuals each harbor a microbiome with different metabolic abilities, personalized responses to different prebiotics have been observed (35, 36). It is therefore important to gain a better understanding of the taxa driving the effects of fructans and other prebiotics on the microbiome to understand how they can impact human health.

Bacterial breakdown and utilization of complex carbohydrates is dependent on the activity of carbohydrate-active enzymes (CAZymes), a highly represented function in the genomes of gut bacteria (10). Glycoside hydrolases (GH) cleave glycans by hydrolyzing glycosidic bonds and represent the largest class of CAZymes. GHs are categorized in families which provide insight on the activity of a particular GH for a specific glycosidic linkage (37, 38). Transcriptional regulators and carbohydrate transporters such as ATP-binding cassette (ABC) and phosphotransferase system (PTS) transporters in the genome are also necessary for carbohydrate utilization (39). In *Bacteroidota* genomes, these CAZymes, transporters, and transcriptional regulators are coregulated and colocalized in polysaccharide utilization loci (PULs) which orchestrate the detection, uptake, and breakdown of glycans (40). While the structure of these genetic loci is not as distinct or well-characterized in non-*Bacteroidota* bacteria, similarly structured gene clusters can be identified in their genomes. Together, the specificities of each PUL element imply the substrate specificity of the PUL, which is an important factor in evaluating which bacteria can metabolize prebiotics of interest (39). However, it is still difficult to predict activity based on PULs or GH families alone, as some families have unknown substrates and some are “polyspecific” or associated with multiple glycosidic linkage activities (39, 41, 42). Furthermore, PULs and GHs in a genome do not always predict activity since they may not be functional or actively used by the bacteria (24, 34).

Multiple approaches have been developed to identify glycan consumers in complex communities such as the gut microbiota. Several recent methods are based on fluorescence-activated cell sorting (FACS) to isolate bacteria from complex communities which interact with glycan probes (33, 43–46). Patnode *et al.*, showed that fluorescently labeled microscopic glass beads containing bound glycans can be used to survey binding specificities of *Bacteroidota* (33). In a similar approach, Riva *et al.* used inulin-grafted nanoparticles to identify inulin-responsive bacteria, with the most responsive bacteria belonging to the *Blautia*, *Collinsella*, and *Faecalibacillus* taxa (43). We recently used different fluorescently tagged glycans to identify and isolate bacterial consumers by FACS. As a result, we were able to identify *Blautia wexlerae* as a novel fructan consumer (44). Together, these methods show that bacteria active on glycans of interest can be identified and isolated, but require the chemical modification or conjugation of glycans, which can represent a limitation.

Metatranscriptomics has been applied to evaluate glycan degradation based on transcriptional activity. This method provides the functional response of a bacterial consortium to glycan supplementation to identify putative consumers (47, 48). Serrano-Gamboa *et al.*, used metatranscriptomics to characterize a consortium of known lignocellulose metabolizers (47). Xu *et al.*, used metatranscriptomics analysis to determine gene expression profiles and microbial abundance in response to raw potato starch, inulin, and pectin supplementation in young pigs (48). Both exploratory methods correlated microbial abundance and CAZyme expression to determine the impact of glycan supplementation in a complex community. However, both methods describe long incubation periods of the consortia with fibres of interest, which can bias bacterial abundances. In Dridi *et al.*, we noticed that induction of metabolic genes resulting in uptake of the fluorescent glycans was in the order of minutes (44). Therefore, we hypothesized that a short incubation time with a glycan would be sufficient to detect the upregulated CAZymes by metatranscriptomics without biasing the bacterial community.

Here, we report a pulse metatranscriptomics approach to identify glycan metabolism genes in a human gut microbiota sample. A fecal sample was incubated for 1 hour in minimum media (MM) or MM containing FOS or galactomannan (GM) as a sole carbon source to induce the expression of glycan utilization genes. To gain a comprehensive understanding of the gut microbial response to glycans, we employed a multi-omics approach integrating metatranscriptomics and whole metagenomic sequencing (WMS). Since GM is an unrelated glycan which requires a separate set of CAZymes, we compared differential abundance of transcripts from mRNA-Seq in samples exposed to FOS versus GM. We found that the upregulated genes were mostly implicated in carbohydrate metabolism with high representation of CAZymes and genes consistent with FOS or GM utilization. We performed WMS on the same stool sample which was then integrated with the mRNA-Seq data to identify the taxa and PULs of potential FOS or GM consumers. Multiple GH32 genes involved in fructan metabolism and overexpressed in the presence of FOS were attributed to *Blautia* species previously unknown to consume FOS. We therefore validated the FOS utilization abilities of several *Blautia* species detected by 16S rRNA sequencing in our sample and confirmed a range of fructan metabolism proficiency. Collectively, our approach takes advantage of the functional response of a gut microbiota to prebiotics to identify and characterize glycan consumers.

## Results

### Brief incubation with glycans induces specific metabolism genes in a complex community

A healthy volunteer donor stool sample was obtained in accordance with the McGill University approved protocol A04-M27-15B and incubated anaerobically for 1 hour in MM or MM supplemented with either FOS or GM as the sole carbon source. Following this short incubation, bacteria from the samples were isolated and their mRNA was extracted to perform mRNA-Seq. We conducted a differential abundance analysis and the predicted protein sequences of the transcripts were annotated using basic local alignment search tool (BLAST) against three protein databases (NCBI nr, UniProtKB Swiss-Prot, and TrEMBL) (Fig 1A).

**Figure 1.**
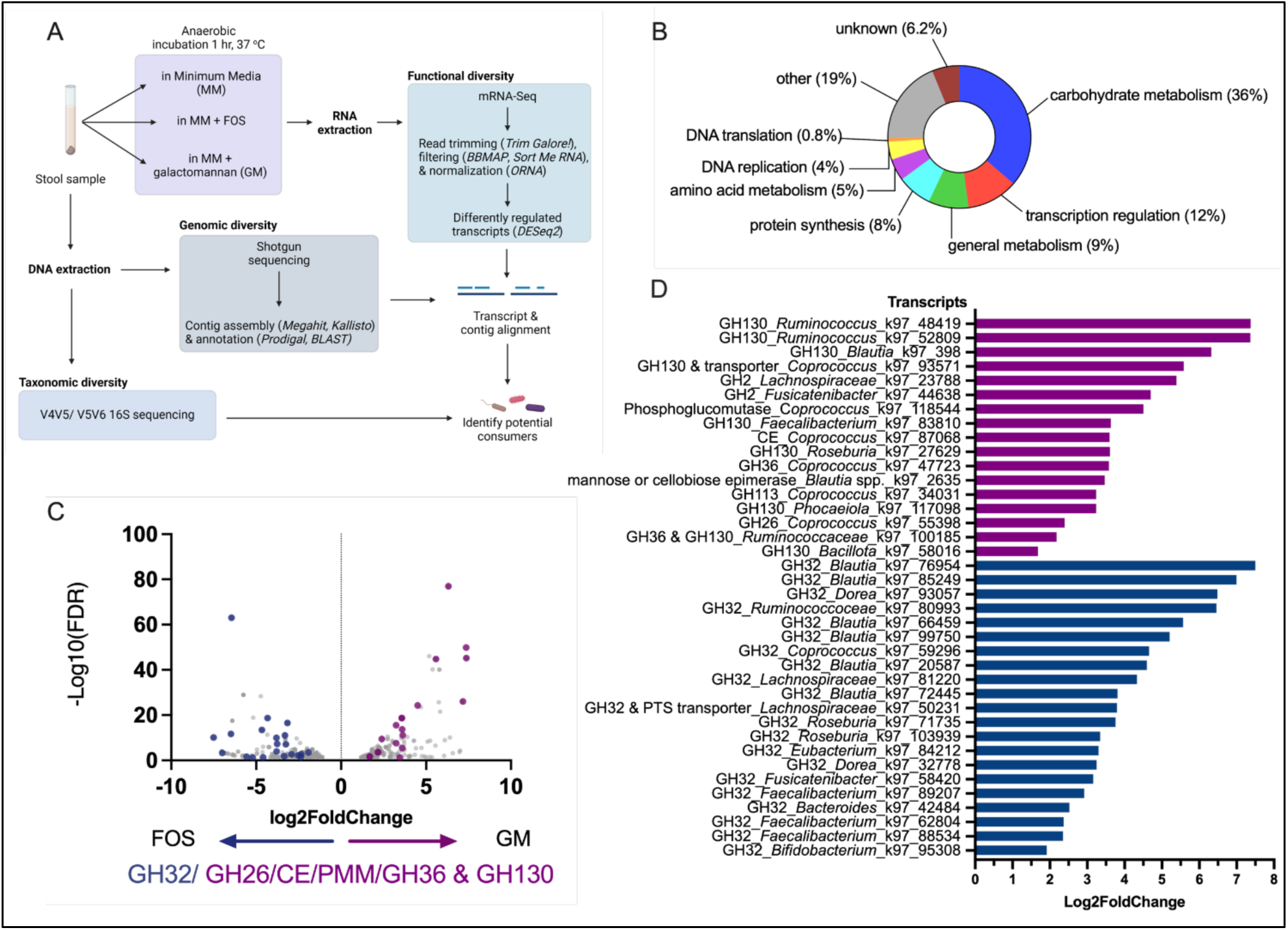
Pulse metatranscriptomics highlights genes upregulated for glycan utilization. (A) Experimental design to identify potential FOS or GM consumers. A stool sample was anaerobically incubated for 1 hour in minimum media (MM) alone or supplemented with FOS or GM as the sole carbon source. Bacterial RNA was then extracted from the culture to perform differential abundance analysis of mRNA transcripts. Bacterial DNA was also extracted from the original stool sample to perform whole metagenomic sequencing (WMS) and 16S rDNA taxonomy identification. (B) Representation of annotated PFAMs belonging to upregulated transcripts. (C, D) DESeq2 differential abundance (Log2Fold Change and false discovery rate (FDR)) of GH-containing transcripts of bacteria cultured in FOS (blue) or GM (purple) expressed as a volcano plot (C) and bar plot (D). An FDR< 0.05, determined by the Benjamini-Hochberg procedure, was considered statistically significant.

We first compared individually the FOS and GM conditions to MM alone and found that in both cases >7,000 genes were differentially expressed. Most induced genes 6,286/7,493 (83.9%) were upregulated in GM (5,980/7,357 (81.3%)) or FOS, compared to MM alone, and most likely represent metabolically active bacteria (Table S1). When comparing samples exposed to FOS vs. GM, we found that only 653 transcripts were differentially abundant (Table S2). Importantly, 36% of the induced genes annotated with a protein domain family (PFAM) were involved in carbohydrate uptake and metabolism, including GHs relevant to FOS and GM utilization (Fig 1B). Moreover, transcription regulation and general metabolism genes represented 12% and 9% of PFAMs, respectively.

Due to the high proportion of metabolism-related genes represented in transcripts upregulated in FOS vs. GM, we used this dataset to further characterize transcripts consistent with FOS or GM utilization. GH32 genes are required to hydrolyze the glycosidic bonds in fructans, and the resultant fructose and glucose monosaccharides are catabolized by bacteria for energy. In contrast, GM oligosaccharides have a β-1,4-D-mannan backbone with branches of D-galactose at the C-6 position (49) that are broken down by GH family GH26, GH36, GH2, and GH130 subfamilies 1 (GH130_1) and 2 (GH130_2) (50, 51). Other enzymes, cellobiose-2-epimerase (CE), phosphomannomutase (PMM), and carbohydrate esterases have also been implicated in GM digestion (51, 52). We therefore searched for transcripts encoding for these genes in the differentially abundant transcripts in both FOS and GM samples (Fig S1). Transcripts containing genes annotated as GH32, also called β-fructofuranosidase or sucrose-6-phosphate hydrolase, were only upregulated in the presence of FOS. In contrast, transcripts containing genes annotated as GH130, GH26, GH36, GH113, CE, β-mannanase, or 4-*O*-β-D-mannosyl-D-glucose-phosphorylase consistent with GM utilization were only upregulated in the samples incubated with GM. We first identified 8 transcripts annotated as CE, PMM, GH26, GH36, GH130, or GH2 and 12 transcripts annotated as GH32s. Since some differentially abundant transcripts lacked an automated annotation, a manual BLASTx was conducted for the transcripts which did not meet our above-mentioned selection criteria. In doing so, we identified 9 additional GM-related genes and 9 additional GH32s. In total, 21 and 17 transcripts encoding genes consistent with FOS and GM metabolism, respectively, were upregulated in their respective conditions (Fig. 1C, D). We also found 6 and 4 transporters or regulators potentially related to FOS and GM utilization, respectively (Fig. S2). Thus, a short exposure with a glycan is sufficient to induce specific metabolic genes expression implicated in FOS and GM metabolism.

### Characterization of GH130 activity

Transcripts consistent with GM metabolism were then identified based on NCBI taxonomy. We found transcripts assigned to taxa previously known as GM consumers such as *Faecalibacterium*, *Ruminococcaceae*, and *Coprococcus* (Fig. 1D) (51–53). For example, *Ruminococcus albus* and *Faecalibacterium prausnitzii* have both been validated extensively for their capability to hydrolyze GMs (51, 54). Of the upregulated transcripts in GM samples (Fig. 1D), 11/17 contained GH130 genes which are active on β-1,2- and β-1,4-mannans (53). To verify the specificity of our identified GH130s on β-1,4-mannans like GM, we used the Sequence Analysis and Clustering of CarboHydrate Active enzymes for Rapid Informed prediction of Specificity (SACCHARIS v2) tool, which allows for the prediction of CAZyme function based on the closest phylogenetic published sequence (Fig. S3) (55, 56). We found that 6/8 selected GH130s were predicted as β-1,4-mannosylglucose phosphorylase (GH130_1; 2.4.1.281) and one was predicted as a β-1,4-mannooligosaccharide phosphorylase (GH130_2; 2.4.1.319) (Fig. S3; Table S3) (51). One transcript was predicted as a β-1,4-mannosyl-*N*-acetyl-glucosamine phosphorylase (2.4.1.320) which is active on *N*-glycans (50). Three sequences were truncated and could not be analyzed SACCHARIS.

### Pulse metatranscriptomics highlight complete *Coprococcus* GM utilization PULs

We found 5 induced transcripts belonging to *Coprococcus*, each encoding a different element of the GM utilization loci. Since *Coprococcus eutactus* has previously been shown to metabolize GM, we wanted to determine if the complete *Coprococcus* GM PUL was represented in our differentially regulated transcripts (52). We therefore conducted WMS of the original stool sample to identify auxiliary genes neighbouring our selected transcripts and evaluate the completeness of the suspected FOS or GM PULs. We integrated the WMS and mRNA-Seq results by aligning the transcripts to the contigs and found that these pairs had ≥98% nucleotide sequence identity. We could map several upregulated transcripts related to GM utilization onto contigs assigned to taxa including *Blautia*, *Ruminococcus*, and *Coprococcus*.

Four of the transcripts from *Coprococcus* putatively involved in GM metabolism were mapped onto contigs associated with a *Coprococcus* using the Clinker pipeline (57), thus reconstructing the GM PUL (Fig. 2). We found that our transcript genes aligned with ≥95% protein homology with reduced homology accounted for by truncation of proteins encoded at the ends of the transcripts. Two transcripts, k97_47723 and k97_93571, aligned alongside each other on to the same contig to form a locus consisting of a *LacI* transcriptional regulator, an ABC transporter, two PMMs, an α-galactosidase (GH36), a 4-O-β-D-mannosyl-D-glycose phosphorylase (GH130_1; RaMP1) and a β-1,4-mannooligosccharide phosphorylase (GH130_2; RaMP2). Transcript k97_87068 containing two cellobiose-2-epimerases (CE) and transcript k97_55398 containing a β-mannanase (GH26) each aligned to separate contigs (Fig 2). This PUL matches the galactomannan PULs extensively characterized by La Rosa *et al.* in *Roseburia intestinalis* and related *Bacillota*, such as *C. eutactus* (52).

**Figure 2.**
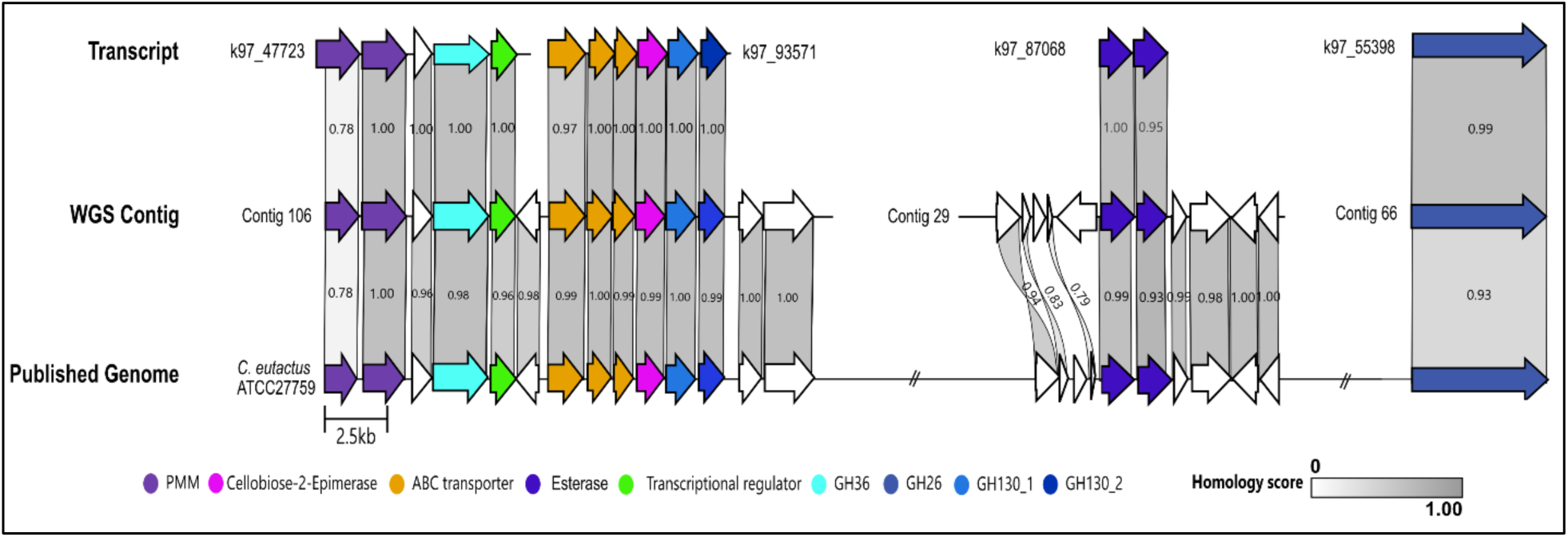
Alignment of predicted galactomannan utilization transcripts to *Coprococcus* WMS contigs and published genome. Upregulated transcripts containing GM-utilization genes were aligned to contigs assigned *Coprococcus* taxonomy and to a published genome for *Coprococcus eutactus* ATCC27759. Using the Clinker pipeline, homologous proteins were aligned and scored with grey bars with a maximum score of 1.00 representing a complete match in protein sequence homology. White arrows indicate unidentified or hypothetical proteins.

To validate our finding, we thus aligned our contigs to the genome of *C. eutactus* (ATCC27759). We found ≥95% protein sequence homology between 19/26 of the genes matched from our contigs and the published genome. However, some genes neighboring the CEs in contig 29 were missing or aligned poorly with 79 - 93% similarity to the neighboring genes in the published genome. The absence or poor representation of these genes in the *C. eutactus* genomes indicates that our contigs belong to another *Coprococcus* species. This is further suggested by the absence of *C. eutactus* in our 16S amplicon sequencing data from that same stool sample (44). Thus, we identified a fully characterized and well established GM PUL in a *Coprococcus* species (52).

### Characterization of GH32 activity and PULs

Transcripts putatively involved in FOS metabolism were similarly found in taxa previously known to be utilizers such as *Faecalibacterium*, *Dorea*, *Eubacterium*, *Roseburia*, and *Coprococcus*, but also *Blautia*, for which only *B. wexlerae* is a validated consumer (Fig. 1D) (24, 44, 58).

GH32s have broad activity across fructans and include enzymes active on both β2-1 linkages found in FOS and inulin, and β2-6 linkages found in levan (59). Therefore, we used SACCHARIS to evaluate the specificity of the GH32 transcripts overrepresented in the FOS samples in our transcriptomics data and found that 18 could be classified as GH32 while 3 selected GH32 transcripts were truncated and could not be classified by SACCHARIS. We found that the vast majority of GH32 sequences (17/18) were most closely related to β-fructofuranosidases, active on β2-1 FOS-type fructans (Fig 3A; Table S4). Only one GH32 (k97_42484) belonging to a *Bacteroides caccae*, was closest to a β-fructofuranosidase, with reported activity on both β2-1 fructans and β2-6 fructans however, the exact substrate specificity is unclear based on our SACCHARIS analysis alone (56, 59).

**Figure 3.**
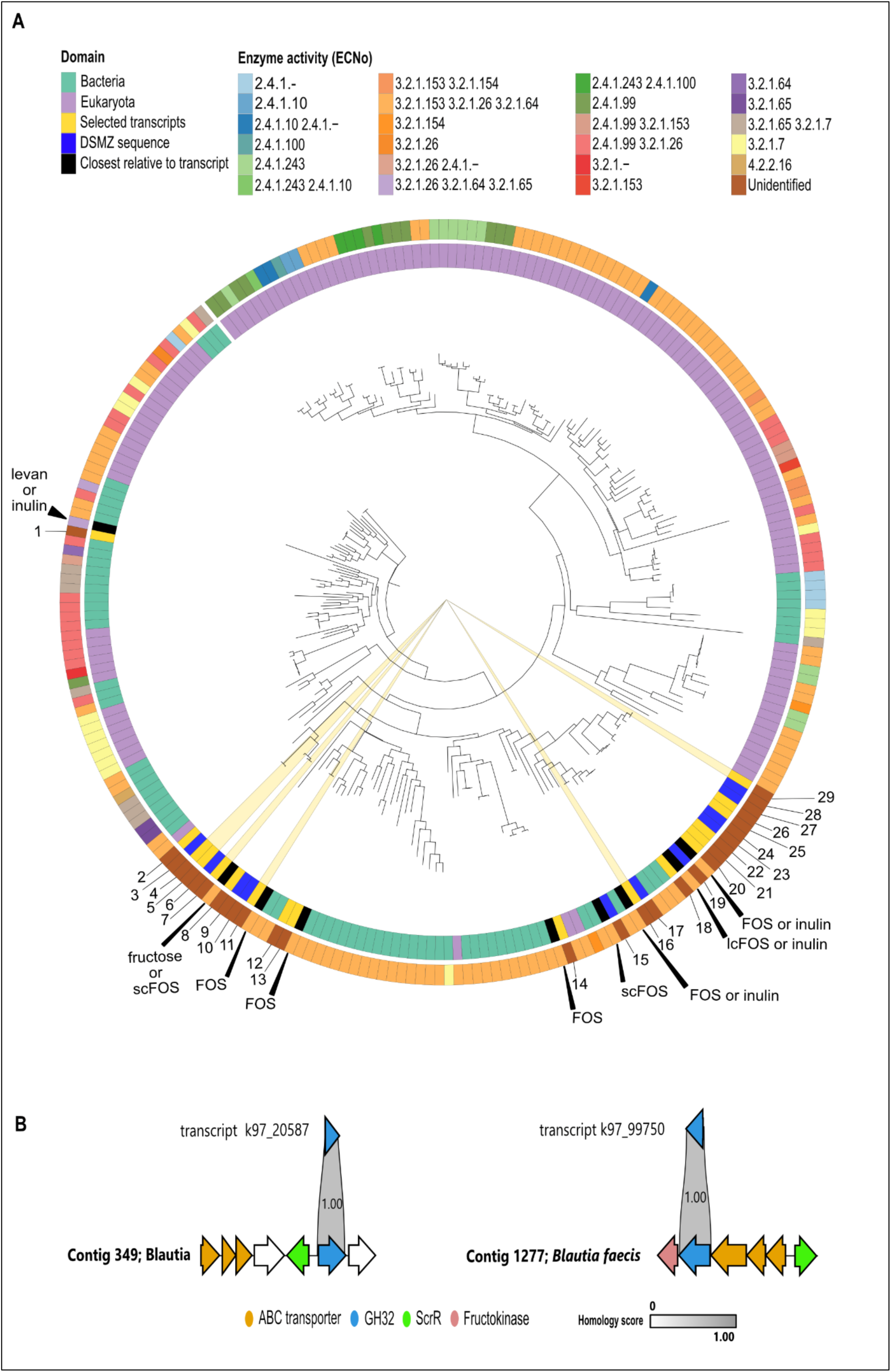
Transcript-contig pairs reveal complete GH32 utilization loci. (A) Phylogenetic tree of GH32 sequences. The phylogenetic trees were generated using the SACCHARIS 2.0 pipeline. The enzyme origin (eukaryotic (purple), bacterial (turquoise), our transcript sequences (yellow), our DSMZ *Blautia* sequences (dark blue), or the closest phylogenetic relative to our transcript (black) is depicted in the inner circle. The enzyme activity category (ECNo) is shown on the outer circle. GH32 transcript sequences assigned to the *Blautia* taxonomy are highlighted by yellow wedges. The closest phylogenetic relatives to each selected transcript are labelled based on their activity on levan, inulin, FOS, or specifically, short chain FOS (scFOS) or long chain FOS (lcFOS). The transcripts are labelled as follows: 1 = k97_42484; 2 = k97_58420; 3 = WP_129257248.1 (*B. faecicola)*; 4 = k97_20587; 5 = k97_85249; 6 = WP_005427754.1 (*B. obeum*); 7 = k97_66459; 8 = k97_103939; 9 = Integrated Microbiome Genomes (IMG) Gene ID 2529070856 (*B. wexlerae*); 10 = WP_059086409.1 (*B. massiliensis*); 11 = k97_76954; 12 = k97_80993; 13 = k97_59296; 14 = k97_95308; 15 = WP_129257123.1 (*B. faecicola*); 16 = k97_72445; 17 = WP_059086463.1 (*B. massiliensis*); 18 = k97_84212; 19 = WP_005425877.1 (*B. obeum*); 20 = k97_88534; 21 = k97_62804; 22 = k97_89207; 23 = WP_154780102.1 (*B. luti*); 24 = IMG Gene ID 2901180188 (*B. faecis*); 25 = k97_71735; 26 = k97_93057; 27 = IMG Gene ID 2529069904 (*B. wexlerae*); 28 = IMG Gene ID 2901180268 (*B. faecis*); 29 = k97_99750. (B) Transcripts k97_20587 and k97_99750 both belonging to a *Blautia* species contain a GH32 (blue arrow), and ABC transporter (orange arrows), and a sucrose operon repressor (*ScrR*; green arrow). Homologous proteins are aligned and scored with grey bars with a maximum score of 1.00 representing a complete match in protein sequence homology. White arrows indicate unidentified or hypothetical proteins.

We were particularly interested in the extent of fructan metabolism in *Blautia,* which is less known, and therefore employed our WMS-transcript integration to characterize the potential *Blautia* fructan PULs. To evaluate the completeness of the PUL surrounding the GH32 in *Blautia* and validate its substrate specificity, we looked at contigs matching those transcripts. Since PTS transporters are usually size-restricted, GH32 genes adjacent to genes encoding a PTS suggests that their substrates are usually shorter DP carbohydrates (phosphorylated-sucrose or -FOSs). In comparison, ABC transporters are associated with longer DP carbohydrates like long-chain FOS or inulin (24). We were able to reconstruct two loci by mapping the GH32 transcripts (k97_20587 and k97_99750) to contigs assigned to *Blautia* species with 100.0% transcript vs contig percent protein homology (Fig. 3B). Both loci contained ABC transporters, suggesting that they are active on FOS or inulin. Furthermore, both transcripts aligned to GH32 loci containing a recombinant sucrose operon repressor (*ScrR*). These *LacI* transcriptional regulators negatively control sucrose transport proteins, further supporting the loci as GH32 PULs (Fig 3B). Additionally, the k97_99750 PUL contains a fructokinase, suggesting it is active on inulin (60–62).

### Fructan utilization varies amongst *Blautia* spp

We then sought to identify the *Blautia* species putatively involved in FOS metabolism in the original stool sample. Based on previously reported V4V5 16S rDNA amplicon sequencing of this stool sample analyzed with the ANCHOR pipeline (63) which can reach species-level identification, five *Blautia* species were found to be present: *B. faecis*, *B. luti*, *B. massiliensis*, *B. obeum*, and *B. wexlerae* (44). We further confirmed the presence of these 5 species by performing V5V6 16S rDNA amplicon sequencing. To investigate fructan utilization by these *Blautia* spp., we used commercially available strains of these species which all had 100% 16S sequence homology in both the V4V5 and V5V6 regions to their representative 16S rDNA sequences from the stool sample (Table S5). In addition, we found that the purchased DSMZ strains encoded GH32s that were clustered closely with the upregulated GH32 transcripts assigned to *Blautia* in a SACCHARIS analysis, suggesting that they likely share substrate specificity (Fig. 3A). In addition, we found a transcript upregulated in the FOS samples (k97_30002) containing ABC transporter genes with >99.3% identity to *B. faecicola* ABC transporters and a *LacI* transcriptional regulator. Therefore, we also obtained this strain, although it was absent in our 16S sequencing.

Ability to metabolize FOS, inulin from chicory, or levan from timothy grass for growth was first assessed by using growth curves of either species in enriched minimal media (MMe) supplemented with each fructan as the sole carbon source (Fig. 4A). Growth for all *Blautia* species, except *B. wexlerae* which we previously assayed, was evaluated in comparison to a glucose or a no-carbon control and quantified based on the difference in area under the curve of MMe alone versus MMe with fructan conditions (Table S6). Four out of 5 strains were able to grow on FOS (Fig. 4A). Metabolism of FOS or inulin varied by species, with *B. faecis* DSM 27629 and *B. massiliensis* DSM 101187 both metabolizing FOS and inulin for growth to levels comparable to glucose conditions. *B. faecicola* DSM 107827 was able to grow in FOS but not to the same extent as with glucose. *B. luti* DSM 14534 also showed some growth in FOS compared to MMe alone, although this growth was significantly lower than in glucose. None of the selected *Blautia* spp. were able to metabolize levan for growth, thus highlighting the specificity of the GH32 subfamilies (Table S6). Growth of *B. obeum* DSM 25238 was inconsistent, possibly because cells quickly lysed, making it difficult to measure culture absorbance (64).

**Figure 4.**
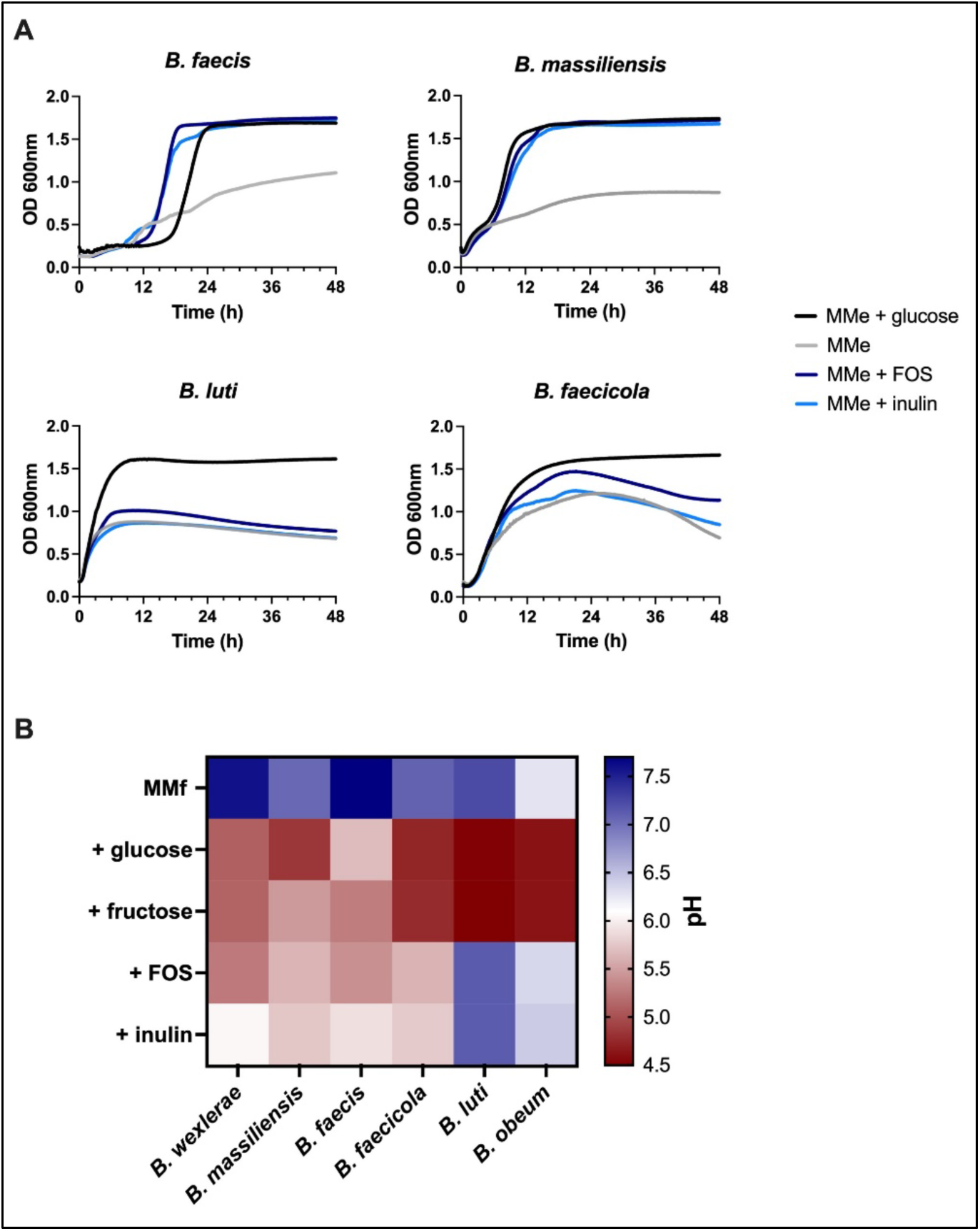
Utilization of fructans varies by *Blautia* species. (A) Growth curves (λ = 600 nm) were conducted for *B. massiliensis*, *B. faecis*, *B. faecicola*, and *B. luti* incubated for 48 hrs in enriched minimal media (MMe; grey) or MMe supplemented with 0.05% FOS (dark blue), 0.05% inulin (light blue), or 0.05% glucose (black) as a positive control. (B) Fermentation assay for *B. wexlerae* (positive control), *B. massiliensis*, *B. faecis*, *B. faecicola*, *B. luti*, and *B. obeum*. Cultures were prepared with 25% BHI (for *B. obeum*) or fermentation minimal media (MMf) supplemented with 1.0% glucose (positive control), fructose, FOS, or inulin. The pH was measured after 48 hrs of incubation to measure pH decrease (red) from the starting pH (blue).

Due to the inconsistent growth of some strains, we also determined acidification of the fermentation minimal media (MMf) by SCFAs to determine the ability of *Blautia* to metabolize fructans (Fig. 4B). As expected from our previous work, *B. wexlerae* DSM 19850 acidified the media in the presence of fructose, FOS, and inulin (44). Mirroring our growth curve results, *B. massiliensis* was able to acidify the media in the presence of all 3 fructans. Similarly, *B. faecis* was able to ferment fructose, FOS, and inulin, as predicted by the complete PUL described by our mRNA-Seq and WMS integration. *B. faecicola* acidified the media in the presence of fructose to a larger extent than FOS or inulin, suggesting that it is more active on shorter chain fructans. Interestingly, *B. luti*, which showed minor growth on FOS based on growth curves, was only able to acidify the media in the presence of fructose. Due to different growth requirements, *B. obeum* was grown in 25% BHI supplemented with 1% glucose, fructose, FOS, or inulin instead of MMf. *B. obeum* was only able to ferment glucose and fructose. These results demonstrate variable fructan metabolism ability by several *Blautia* strains in addition to *B. wexlerae*, with *B. faecis* and *B. massiliensis* being particularly active on FOS and inulin.

## Discussion

Glycans are widespread in human diet and play a pivotal role in shaping the gut microbiota and its effect on host health. Galactomannans are ubiquitous in modern human diet as the food stabilizer guar gum or locust bean gum and fructans now feature as key ingredients in products promoting gut health benefits (58, 65). As glycan-based prebiotics become a promising therapeutic option for a variety of conditions, it is important to identify which members of the gut microbiota can metabolize glycans of interest and the genes and pathways involved (22, 23, 31, 31, 66). By linking bacterial metabolic genes to their substrates, we can gain a better understanding of the mechanistic link between dietary fibre and its health benefit. Since the presence of GHs alone is insufficient to evaluate a microbiome’s potential to breakdown glycans, there is a need to further contextualize GH activity in response to prebiotics. Identifying and validating GHs in the gut microbiome will also help to improve our prediction of GH specificity by bioinformatic tools such as SACCHARIS. As prebiotic responses vary amongst individuals, an increased catalog of characterized glycan metabolism genes will allow us to predict the prebiotic fermentation potential of a given individual, according to their unique microbiome.

Here, we demonstrate that pulse metatranscriptomics, which capitalizes on the rapid induction of carbohydrate metabolism-related genes, integrated with WMS, can identify genes and bacteria responsible for glycan metabolism in a human stool sample. Analysis of gene induction in MM compared to FOS or GM conditions revealed several differentially regulated genes, many of which were unrelated, general metabolism genes or were genes pertaining to functions in metabolically active bacteria. To isolate PUL-based metabolism of complex glycans, other approaches have used glucose catabolic repression which may be a promising alternative to incubation in MM alone (67). Here, we compared gene induction in FOS versus GM conditions directly, since they require distinct metabolic genes. We noted active induction of FOS utilization genes by previously described consumers including *Bacteroides*, *Coprococcus*, *Dorea*, *Faecalibacterium*, and *Roseburia* (24, 58, 59). We also saw induction of GM utilization genes in *Blautia*, *Coprococcus*, and *Ruminococcus,* also previously shown to metabolize GM (52, 68–71). This is not surprising since both glycans have been well studied, with the CAZymes required for the hydrolysis of either glycan described in several bacterial taxa (24, 51, 52, 59). Importantly, we only saw the induction of putative CAZymes in the presence of their respective substrate. Furthermore, the integration of transcriptomics and WMS data from the same stool sample allowed us to reconstruct GM utilization gene loci consisting of 14 genes spread across 4 PULs from a *Coprococcus* species. These GM loci matched the validated GM utilization PULs characterized in *C. eutactus* (52).

Importantly, despite FOS being a well-studied glycan, we identified several members of the *Blautia* genus as novel consumers. *Blautia* has previously been shown to increase in abundance in the presence of inulin or FOS *in vitro* in human stool and *in vivo* in rats and mice (15, 68, 71, 72). *Blautia* are amongst the dominant genera of the human intestine and were first isolated from human stool (73). Most interestingly, increased abundance of *Blautia* has been negatively correlated with visceral diabetes, anti-lipidemia, and obesity-associated inflammation (74–78). *Blautia wexlerae* is also correlated with a favourable response to immune checkpoint inhibitors in multiple cancer patient cohorts (79). In addition, a high abundance of *Blautia* as well as supplementation of *Blautia coccoides* in humanized mice has been shown to produce SCFAs from a fibre rich diet which can stimulate basal mucosal secretion and maintain colonic mucus (80). Increasing baseline fibre intake rather than supplementing *Blautia* probiotics alone may be a promising alternative to stimulate mucus growth in the colon and prevent defective mucus function which has been linked to metabolic diseases and obesity (80, 81). For these reasons, it is important to gain a better understanding of how gut bacteria, like *Blautia*, breakdown dietary fibre and in turn, impact host health.

We found that *B. massiliensis*, *B. faecis*, and *B. faecicola* were clear metabolizers of both FOS and inulin; yet fructan metabolism was limited in *B. luti* and *B. obeum*. With our growth curve assays, we saw minor *B. luti* growth in the presence of FOS but no fermentation. Similarly, *B. obeum* could neither ferment nor utilize FOS for growth; yet both *B. luti* and *B. obeum* can ferment fructose. These results demonstrate that glycan utilization varies at the species level and potentially, as has been previously shown, at the strain level (33). This also suggests that the GH32 present in the *B. luti* and *B. obeum* genomes are active on fructose of FOS with low DP (24). Growth or fermentation observed in our assays can likely be attributed to the presence of such low DP fructan in the commercially available FOS used. The *B. obeum* or *B. luti* GH32s in our mRNA-Seq dataset could also have been activated by the presence of fructose or low DP FOS produced by other members of the bacterial community. In many cases, this cross-feeding can structure how prebiotics are broken down by the gut microbiota (82). Bacteria which initially breakdown a polysaccharide (primary consumer) and produce oligosaccharides of lower DP which become accessible to other bacteria for consumption (secondary consumers), which makes for more efficient glycan breakdown (9, 83). For example, the degree of FOS or inulin breakdown in a microbiota can be dictated by the ratio of *Bacteroidota*/*Bifidobacteria* in the community, as these taxa have an established cross-feeding relationship (84).

The use of commercial *Blautia* isolates is a limitation to our method in determining fructan utilization in *Blautia* spp.. Indeed, strain differences between the purchased strains and the strains present in our stool sample may explain differences in fructan utilization as CAZyme presence and substrate affinity varies at both the species and strain levels (10). However, SACCHARIS clustering suggests that the commercial strains are close relatives to the *Blautia* in the sample in terms of GH32 homology. Additionally, some cases of fructan utilization may have been missed in certain species because of the lack of co-factors or primary degraders in the cultures of the commercially available *Blautia* strains (82). Together, these results suggest that bacteria can identify and respond to the presence of glycans which they may not be equipped to metabolize (34, 44). While this is a limitation, the strength of our method lies in its specificity towards the glycans of interest. Furthermore, taxa identification in transcriptomics is typically restricted to the genus level. Here, we utilized 16S sequencing data from the same stool sample to enhance our understanding of the microbial community composition. In the future, complementing pulse metatranscriptomics with established methods which couple metabolic labeling to FACS such as in Dridi *et al.*, might allow isolation of glycan metabolizing strains from the stool sample (44).

Here, we identified several *Blautia* species as novel consumers of FOS, a thoroughly studied prebiotic. Applying this method to lesser characterized glycans or other dietary components, will help identify putative consumers of prebiotics which may be applicable in health conditions where the gut microbiome is implicated. This method allows us to characterize genes involved in prebiotic metabolism, potentially leading to the discovery of novel CAZymes or PUL assemblies, especially in non-*Bacteroidota* bacteria. *Bacillota* are recognized as key prebiotic consumers and prolific SCFA producers, known to deploy elaborate systems for polysaccharide utilization (85). The discovery of non-*Bacteroidota* consumers via pulse metatranscriptomics can broaden the scope of glycan utilization strategies across species. As we continue to untangle the relationship between the gut microbiota and host health, understanding the impact of diet on gut commensals becomes increasingly important. At a larger scale, this knowledge will allow us to determine the individual response to dietary fibre by each person’s unique microbiota, allowing the development of targeted prebiotic approaches to provide the desired effect on host health.

## Acknowledgments

This research was funded by a Canadian Glycomics Network (GlycoNet) Catalyst Grant (DO-16) to B.C. and C.F.M. and by the Canadian Institutes of Health Research (CIHR) grant PJT-437944 to B.C. and C.F.M. B.C. holds a tier II Canada Research Chair (CRC) in Therapeutic Chemistry. C.F.M. holds a tier II CRC in Gut Microbial Interactions. The sequencing was performed at the McGill University and Genome Quebec Innovation Centre.

## Methods

### Stool incubation and RNA isolation

All preparation and incubations were performed under anaerobic conditions (85% N_2_, 10% CO_2_, and 5% H_2_). A 0.3 g fecal sample was weighed and diluted to 1:10 g/mL in minimum media (MM) (6.6 mM KH_2_PO_4_ (pH 7.2), 15 mM NaCl, 100 μM MgCl_2_, 175 μM CaCl_2_, 50 μM MnSO_4_, 5 mM (NH_4_)_2_SO_4_, 15 μM FeSO_4_, 24 μM NaHCO_3_, 1 g/L L-cysteine, 1.9 μM hematin, 6 μM hemin, and 200 ng/ml−1 vitamin B_12_), vortexed thoroughly and centrifuged for 3 min at 700 × *g*. The supernatant was saved, and the pellet was discarded. The supernatant was centrifuged for 5 min at 6,500 × *g*, the supernatant was discarded, and the pellet was washed with MM. The final pellet was resuspended in MM supplemented with FOS (FOS from chicory, F8052, Sigma, Canada) or GM (low viscosity carob galactommanan, P-GALML, Neo Gen, USA) at 0.1%. Control was realized by the incubation of the pellet in MM. The bacteria were incubated at 37 ᵒC for one hour. After incubation, the bacteria were collected by centrifugation 6,500 x *g* for 5 min and resuspended in 200 µL of MM prior to RNA extraction. RNA isolation was then performed using the AllPrep PowerFecal DNA/RNA kit (Qiagen, Canada) according to the manufacturer’s instructions. A DNA digestion step was performed on the isolated RNA using the Ambion DNA-free DNA Removal Kit (Invitrogen) according to the manufacturer’s instructions, to ensure complete DNA removal. From the same stool (MX73), three replicates for each bacterial incubation were performed. For each bacterial incubation, the RNA extractions were performed in triplicate and pooled.

### RNA sequencing

mRNA-Seq was performed by Génome Québec. In summary, total RNA was prepared for Illumina sequencing using the NEBNext rRNA Depletion Kit to remove rRNA and using the NEBNext Multiplex Oligos for Illumina. Prepared libraries were quality checked with a Bioanalyzer 2100 (Agilent) prior to sequencing. Sequencing was performed on a HiSeq 4000 (Illumina), Paired End 100bp, sequencing lane 300 M reads, generating an average of 30 M reads per sample (20.35–36.47 M).

### Stool DNA extraction and shotgun sequencing

As previously described in Dridi et al., total bacterial DNA from stools was extracted using the AllPrep PowerFecal DNA/RNA kit (Qiagen, Canada) following the manufacturer’s instructions. DNAs were quantified by the Qubit Fluorometric Quantification method (Invitrogen). DNA was sent to Génome Québec for Illumina HiSeq 4000 PE 100 bp sequencing for shotgun metagenomics (44).

### 16S ribosomal RNA gene amplification, and sequencing

DNA was extracted from the stool sample using the AllPrep Powerfecal DNA/RNA kit (Qiagen, Canada) following the manufacturer’s instructions. DNA was quantified by the Qubit Fluorometric Quantification method (Invitrogen). The primers GGMTTAGATACCCBDGTA (F) and GGGTYKCGCTCGTTR (R) were used to target the V5V6 region. The CS1 (ACACTGACGACATGGTTCTACA) and CS2 (TACGGTAGCAGAGACTTGGTCT) tags were used to add barcodes and Illumina adapters. The Q5 High Fidelity DNA polymerase (New England BioLabs) was used to perform amplification with the PCR cycles starting with an initial denaturation step of 98 ᵒC for 30 seconds followed by 23 cycles of: (1) 98 ᵒC for 10 s; (2) 58 ᵒC for 15 s; (3) 72 ᵒC for 30 s, with a final extension at 72 ᵒC for 2 min. The MiSeq platform was used for 2 x 250 bp paired-end sequencing of the resulting PCR products. Sequence data was deposited is available under the BioProject PRJNA925842.

### RNA-Seq / WMS assembly and annotation

Trim Galore! (v0.6.6) (86), a wrapper script utilizing Cutadapt and fastQC (87) was used with recommended parameters for adapter and quality trimming as described in the manual. BBMAP (v.38.90) (88) removed potential human contamination using the masked hg19 human assembly. Reads originating from rRNAs were filtered out using SortMeRNA (v2.1) (89). ORNA (v1.0) (90) normalized read data.

MEGAHIT (v1.2.9) (91) was used to assemble reads from all samples into one co-assembly using meta-large option. Kallisto (v0.46.2) (92) assigned reads and inferred contig/transcript abundance using expectation maximization (93). Prodigal (v2.6.3) (94) predicted open reading frames (ORFs) with the “meta” option, and BLAST (95) annotated contig sequences.

To assign contig/transcript taxonomy full contig lengths were aligned against the NCBI nt database and Reference Viral Database (RVDB v25.0) using BLASTn with the following parameters: -evalue 1e-50, -word_size 128, and -perc_identity 97.

For gene annotation, translated predicted protein sequences were initially searched against three protein databases (NCBI nr, UniProtKB Swiss-Prot, and TrEMBL) using BLASTx with the following parameters: -evalue 1e-10, -word_size 6, and -threshold 21. The best bitscore from each database was selected for each ORF. Subsequent annotations were derived by mapping Gene Ontology (GO), Pfam, PANTHER, EMBL, InterPro, HAMAP, TIGRFAMs, STRING, HOGENOM, and SUPFAM terms from the UniProtKB database. To further supplement functional and pathway information, amino acid sequences were submitted to the GhostKOALA webserver (96). KEGG functional and taxonomic annotation was then retrieved using both complete and incomplete pathways.

### Differential abundance/expression analysis

Prior to differential abundance analysis, sparsity and count thresholds were applied, requiring that a contig/transcript count in a single sample be <90% of the count across all samples and that contig/transcript occurrence be at least ≥3 in samples within the same design factor. Differential abundance (or expression) analysis was performed using DESeq2 (97) on pre-processed raw abundance data of contigs/transcripts. Normalization and variance stabilization were achieved using the regularized logarithm (rlog) transformation. A false discovery rate (FDR) < 0.05, determined by the Benjamini-Hochberg procedure, was considered statistically significant.

### Phylogenetic analysis of GH32 and GH130 enzymes

Transcript sequences were submitted as input to SACCHARIS v2 for phylogenetic analyses (55, 56). Briefly, sequences and accession numbers of characterized GH32 and GH130 enzymes were extracted from the CAZy database (July 2024) (38) and, with transcript sequences, were pruned to the respective GH domains as identified by dbCAN2 (98). ModelTest-NG (99) was used for best-fit model selection using the sequence alignment, and FastTree (100) was used to generate the phylogenetic trees. Trees were then annotated using the RSACCHARIS package (55) in RStudio (101) with R version (4.3.2).

### PUL assembly

Selected contigs were annotated with Rapid Annotation using Subsystem Technology (RAST) (102). The annotated contigs and transcripts were then aligned with CompArative GEne Cluster Analysis (CAGECAT) Clinker pipeline (57).

### Bacterial strains and pre-culturing

All *Blautia* species were purchased from DSMZ (strains are listed in Table S5) and grown at 37 °C in an anaerobic chamber (5% H_2_, 20% CO_2_, 75% N_2_; Coy Lab Products). *Blautia wexlerae* (strain DSM 19850) was grown on Columbia agar 5% plates (Northwest Scientific, Inc.) and all other *Blautia* strains were grown on brain-heart-infusion plates (Oxoid). A colony of each bacterium was inoculated into fresh brain-heart-infusion broth (BHI; Oxoid) and cultured at 37 °C for 24 or 48 h prior to the start of growth experiments.

### Growth curves

A modified minimal media (MMe; 6.6 mM KH_2_PO_4_ (pH 7.2), 15 mM NaCl, 100 μM MgCl_2_, 175 μM CaCl_2_, 50 μM MnSO_4_, 5 mM (NH_4_)_2_SO_4_, 15 μM FeSO_4_, 24 μM NaHCO_3_, 1 g/L L-cysteine, 1.9 μM hematin, 6 μM hemin, and 200 ng/ml−1 vitamin B_12_, 7 mg/mL yeast extract, 3.5 mg beef extract. Filter sterilized with a 0.2 µl Filtropur S syringe filter (Sarstedt Inc)) supplemented with 0.05% glucose (Sigma G8270-100G), 0.05% FOS (Sigma F8052-50G), 0.05% inulin (Sigma I2255-10G), or 0.05% levan (Megazyme P-LEVAN) was prepared and filter sterilized (0.2 µm pore size; Sarstedt). In a 96-well plate (Corning), 245 µl of each enriched minimal media was added per well in triplicates and inoculated with 5 µl of preculture. Growth curves were performed anaerobically in a plate reader (Epoch) at 37 °C, and in cycles of 10 minutes, the plates were orbitally shaken for 10 seconds before OD-measurement at 600 nm (OD600) over a period of 48 h at which points cultures reach stationary phase. Growth curves were then calculated based on the average of the OD600 triplicate values. The Δ_Area under the curve_ (ΔAUC) was calculated and growth on 0.05% FOS, 0.05% inulin, or 0.05% levan was quantified in comparison to growth in the positive control (0.05% glucose supplemented MM). Dunnet test was performed with glucose as a control to compare growth in different media conditions.

### Fermentation assays

Fermentation minimal media (MMf; 6.6 mM KH_2_PO_4_ (pH 7.2), 15 mM NaCl, 100 μM MgCl_2_, 175 μM CaCl_2_, 50 μM MnSO_4_, 5 mM (NH_4_)_2_SO_4_, 15 μM FeSO_4_, 24 μM NaHCO_3_, 1 g/L L-cysteine, 1.9 μM hematin, 6 μM hemin, and 200 ng/ml−1 vitamin B_12_, 1 mg/mL yeast extract, 0.5 mg/mL mg beef extract, 1 mg/mL casitone, 1 mg/mL soyatone, 0.025 mg/mL bromoscerol purple (Sigma 114375-5G). Filter sterilized with a 0.2 µl Filtropur S syringe filter (Sarstedt Inc)) was supplemented with 0.05% glucose (Sigma G8270-100G), 0.05% fructose (Sigma F0127-100G), 0.05% FOS (Sigma F8052-50G), or 0.05% inulin (Sigma I2255-10G). *B. obeum* was grown in 25% BHI (Oxoid) supplemented with 1.0% glucose, 1.0% fructose, 1.0% FOS, 1.0% inulin. The acidification assay was performed by inoculating 2 ml of glycan-supplemented MMf with 50 µl of preculture followed by anaerobic culturing for 48 h (this was performed in triplicates). In the case of *Blautia faecis* (strain DSM 27629) and *Blautia faecicola* (strain DSM107827), the incubation period was extended to 92 h. Glycan metabolism was determined by acidification based on pH values recorded with a pH meter.

### Data availability

Sequence data was deposited in GenBank (Sequence Read Archive) and is available under the BioProject PRJNA925842 (https://www.ncbi.nlm.nih.gov/bioproject/?term=PRJNA925842). Sequencing of the V4V5 16S rRNA sequencing for this sample (MX73) was previously described and published (44).

## Supplementary figures

**Figure S1.**
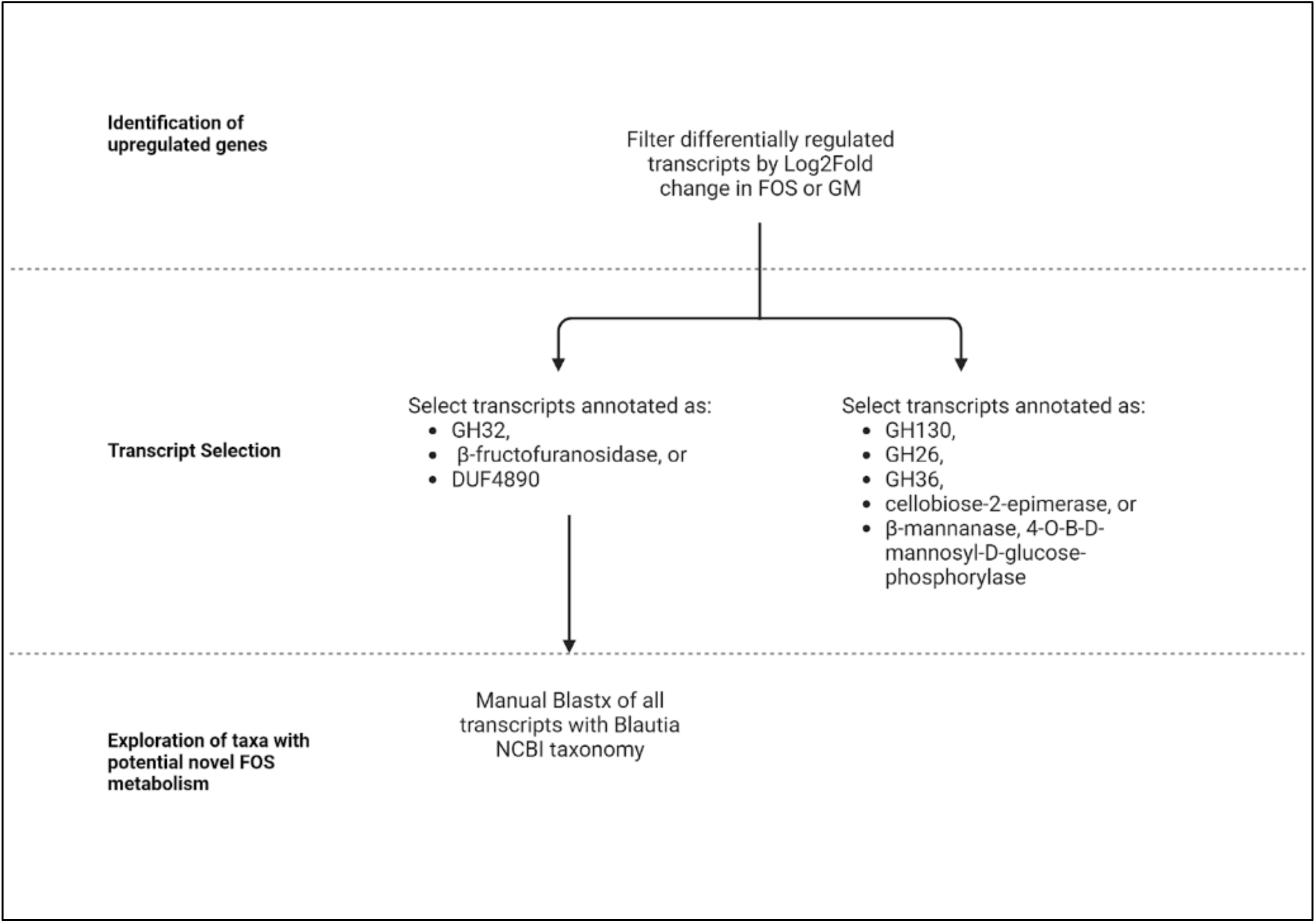
Workflow of RNAseq transcript selection process.

**Figure S2.**
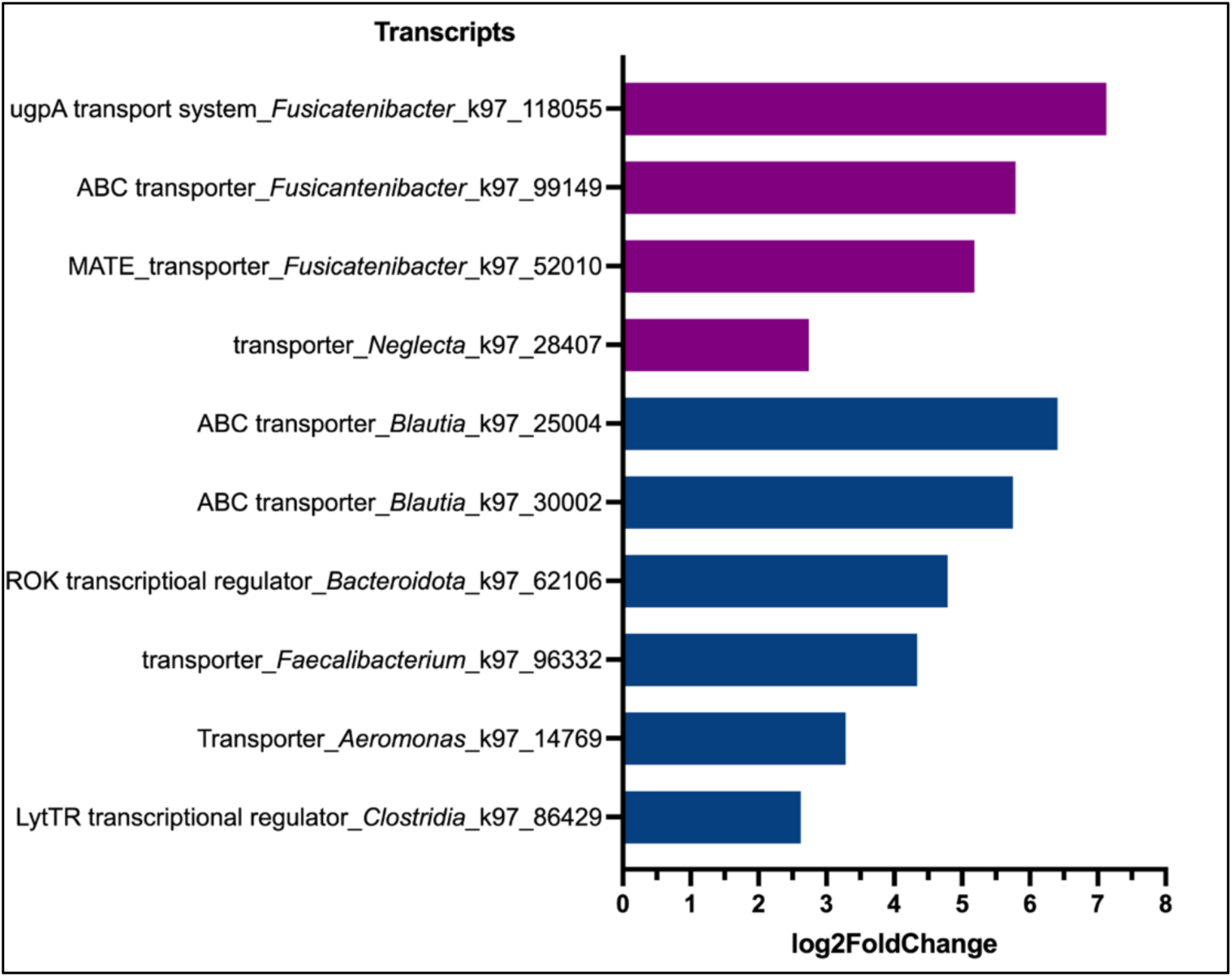
RNASeq differential abundance of transcripts containing transporters or transcriptional regulators related to FOS (blue) or galactomannan (purple) metabolism.

**Figure S3.**
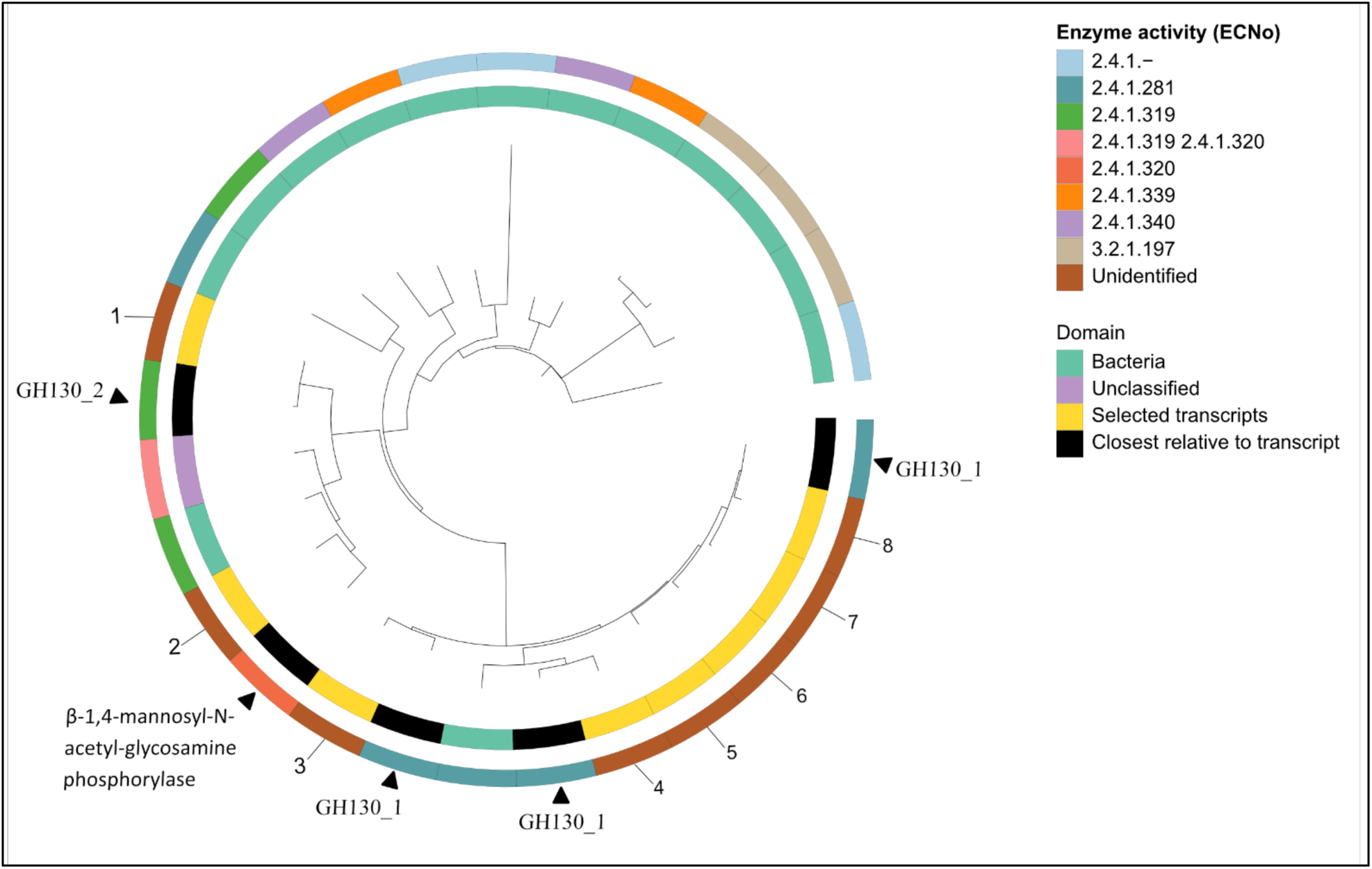
SACCHARIS analysis of GH130s substrate affinity. The phylogenetic trees were generated using the SACCHARIS 2.0 pipeline. The enzyme origin (bacteria (turquoise), unclassified (purple), or our transcript sequences (yellow)) is depicted in the inner circle. The closest neighbouring sequences are depicted in black and are active on β-mannans ((GH130 1; β-1,4-mannosylglucose phosphorylase; 2.4.1.281), (GH130_2; β-1,4-manooligosaccharide phosphorylase; 2.4.1.319)) or on N-glycans (β-1,4-mannosyl-N-acetyl-glucosamine phosphorylase; 2.4.1.320). The enzyme activity (ECNo) is shown in the outer circle.

**Table S5.** Strains used in this study.

| Strain | Source | Distributer | % Identity to ANCHOR V4V5 | % Identity to ANCHOR V5V6 |
| --- | --- | --- | --- | --- |
| *Blautia faecicola* DSM 107827 | Human faeces (103) | DSMZ GmbH | NA | NA |
| *Blautia faecis* DSM 27629 | Human faeces (104) | DSMZ GmbH | 100.0 | 100.0 |
| *Blautia luti* DSM 14534 | Human faeces (105) | DSMZ GmbH | 100.0 | 100.0 |
| *Blautia massiliensis* DSM 101187 | Human faeces (106) | DSMZ GmbH | 100.0 | 100.0 |
| *Blautia obeum* DSM 25238 | Human faeces (107) | DSMZ GmbH | 100.0 | 100.0 |
| *Blautia wexlerae* DSM 19850 | Human faeces (105) | DSMZ GmbH | 100.0 | 100.0 |

**Table S6.** Difference in area under the curve (ΔAUC) between MMe alone growth curve and each growth curve of MMe supplemented with 0.05% glucose, 0.05% FOS, 0.05% inulin, and 0.05% levan. Dunnet test was performed with glucose as a control to compare growth in different media conditions. ** indicate p<0.005; *** indicate p<0.0005

| Blautia species | Enriched Minimum Media |  |  |  |
| --- | --- | --- | --- | --- |
|  | + glucose | + FOS | + inulin | + levan |
| *faecicola* | + 22.03 | +10.60<br>($p = 0.1121$) | +1.87<br>(** $p = 0.0085$) | -2.09<br>(** $p = 0.0030$) |
| *faecis* | +17.70 | +24.84<br>($p = 0.2720$) | +1.85<br>($p = 0.4251$) | -3.14<br>(** $p < 0.0027$) |
| *luti* | +34.85 | +4.40<br>(*** $p < 0.0001$) | -0.44<br>(*** $p < 0.0001$) | -0.26<br>(*** $p < 0.0001$) |
| *massiliensis* | +34.72 | +32.95<br>($p = 0.7819$) | +30.93<br>($p = 0.1702$) | -1.93 ***<br>($p < 0.0001$) |

